# Chromosome compartmentalization replacement during stem cell differentiation

**DOI:** 10.1101/284190

**Authors:** Y.A. Eidelman, S.V. Slanina, V.S. Pyatenko, S.G. Andreev

**Affiliations:** Institute of Biochemical Physics, Russian Academy of Sciences, Moscow, Russia; A.Tsyb Medical Radiological Research Center, branch of the National Medical Research Radiological Center of the Ministry of Health, Obninsk, Russia; National Research Nuclear University MEPhI, Moscow, Russia

## Abstract

In this paper, changes in a large-scale 3D structure of chromosomes during stem cell differentiation is studied. The polymer coarse-grained model of a human interphase chromosome is introduced which reproduces the experimental Hi-C contact maps in chromosomes 12, 17 for both embryonic stem and differentiated cells with high accuracy. Model based analysis of Hi-C data suggests a mechanism of establishment of preferential long-range chromosomal contacts and compartmentalization replacement during cell stem differentiation. The model provides the conceptual basis for integration of data on the dynamics of chromatin interactions, the 3D structure of chromosomes and gene expression during stem cell differentiation or reprogramming.

## INTRODUCTION

Folding of chromatin in the nucleus is thought as the important factors in the regulation of gene expression [1, 2]. But whether this reflects a general genome principle is still unknown. Advanced approaches to the 3D genome study use ligation of spatially proximal loci and are called “3C-based technologies”, such as 3C, 4C, 5C and Hi-C [3–6]. The development of 3C-based technologies, together with high-performance sequencing, allowed to integrate large amounts of data and to obtain the new information on spatial genome architecture [7, 8]. A multi-level organization of chromatin has been shown, from (sub)megabase chromatin domains, or topologically associating domains (TADs) [9–11] to the supra-megabase level where the complex, “checkerboard” contact maps has been observed [6, 12, 13]. The checkerboard structure of contact maps is often interpreted as follows: the entire genome (or each chromosome) consists of two different types of compartments, A and B, so that domains belonging to the same-type compartments contact preferentially and those belonging to different-type compartments contact rarely [6]. Belonging to compartments A/B is correlated with various genomic properties, including GC content, DNAse-I hypersensitivity sites, histone methylation and acetylation, replication timing, *etc*. [2, 6, 14]. Compartment A is associated mainly with active chromatin, compartment B with inactive chromatin [15].

During differentiation, change of expression levels is accompanied by coordinated switching between A and B compartments [16]. Intensive interactions between TADs have been observed and the concept of meta-TADs has been introduced [17]. Changes in transcription during development have been shown to correlate with chromatin insulation, the establishment of boundaries of chromatin domains [18]. During mouse neuronal differentiation, contacts between active TADs become less pronounced, while inactive TADs contact more frequently [18]. These data are consistent with early proposals linking genome 3D organization with stages of differentiation [19, 20]. These ideas are in concert with explanation of Waddington landscape [21] in terms of higher-order chromatin organization [20].

There are numerous gaps in understanding the mechanisms of how chromosomes fold in a highly condensed nuclear environment. In order to gain insight into folding mechanisms, methods of bioinformatics and polymer physics are widely used for retrieving relevant information from Hi-C data. Polymer approaches are considered to be most adequate in obtaining characteristics of chromatin and chromosomes with taking into account their polymeric nature [6, 11, 22–26]. The difficulties of physical modeling of chromosomes do not allow yet to find comprehensive answers to many questions, in particular, quantitative explanation of chromosomal contact heatmaps remain not completely understood [27–30].

In this paper, we study the 3D organization of chromosomes during stem cell differentiation. A physical model of human interphase chromosomes 12 and 17 is developed and experimental Hi-C data for embryonic stem (hESC) and differentiated (IMR90 fibroblasts) cells are analyzed. The model reproduces the observed contact matrixes for chromosomes 12 and 17 in both cell types with high accuracy, thus demonstrating ability to solve the problem of a high-precision description of chromosomal contacts. The results reveal that the transition from the pluripotent to differentiated state is characterized on the chromosomal level by differential change of interactions between chromosomal domains in various chromosomal regions. Model based analysis of Hi-C data suggests a mechanism for the establishment of long-range contacts preferential for given cell type. In such mechanism the transition from pluripotent to differentiated state is triggered by reorganization of chromatin architecture. Rearrangements of the chromosome topology, shaped by a network of volumetric interactions, can serve as driving force for compartmentalization replacement during cell stem differentiation or its reversal during dedifferentiation.

## MATERIALS AND METHODS

The structure of the interphase human chromosome 12 is modeled by a non-lattice model in the form of a flexible heteropolymer chain of semipermeable spherical subunits with a DNA content 100 thousand base pairs, kbp. Various types of coarse-graining (and approximation) are typically used in modeling chromosomes [31]. The combined structure of the chromosome is based on the concept of organizing chromosomes from megabase domains or Topologically Associating Domains (TADs) [9, 11]. Domains, or TADs, have a complex internal structure [10, 11], but due to computational limitations we consider TADs as structureless subunits, ignoring simultaneous calculation of intra- and inter-TAD interactions, and for analyzing full-chromosome maps with resolution 1 million base pairs (Mbp) we limit ourselves to considering interactions between 1 Mbp chromosomal blocks, which roughly corresponds to TAD-TAD interactions. In the calculations for particular chromosomes, the number of subunits is determined on the basis of their length in Mbp, i.e. chromosome 12 consists of 1350 structural 100kbp subunits, or monomers, chromosome 17 consists of 820 monomers. Conformational changes and spatial packing of chromosomes in the nucleus space are the result of the interaction of remote subunits, which (in the physics of polymers) are called volumetric interactions [32, 33].

The potential of volumetric interactions between a pair of elements (*i, j*) takes into account two components, the excluded volume and the attracting part, *U*_*ij*_(*r*) = *U*_*ev*_(*r*) + *U*_*attr*_(*r, i, j*), where *U*_*ev*_(*r*) is the potential of excluded volume, i.e. repulsion of elements at short distances, *U*_*attr*_(*r, i, j*) is an attractive potential taking into account in the generalized form heterogeneous DNA-protein interactions, intranuclear medium (ionic strength, pH), *etc*. The potential of the excluded volume in this model is the shifted Lennard-Jones potential:

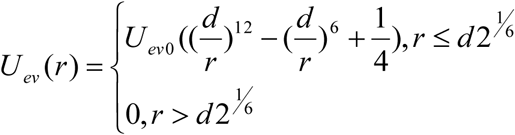

where *r* is the distance between the centers of two elements, *d* is the diameter of the element. The value of *U*_*ev0*_ does not depend on the position of the elements on the chromosome, i.e. on the numbers *i* and *j*. The meaning of the shifted potential is that it is continuous, because it vanishes at *r* = *d*·2^1/6^. The attracting potentials are cut and shifted Lennard-Jones potentials with coefficients depending on the position of the elements along the chromosome:

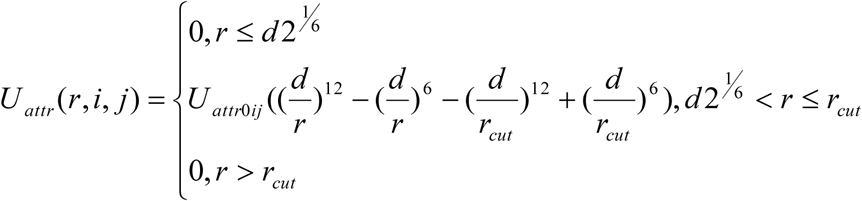

where *r*_*cut*_ is the distance between the centers of the elements, from which the potential is not calculated. This is a standard technique for reducing computational load. We use the value *r*_*cut*_ = 3*d*, at which the non-displaced value of the potential is 0.5% of the depth of the well and therefore we can neglect attraction at distances greater than *r*_*cut*_.

The idea of a heteropolymer derives from the modeling of TADs, where all potentials were considered different [34, 35]. We generalize this idea to the level of the whole chromosome. In contrast to our previous works [28, 29], where the Monte Carlo technique was used, here we use the molecular dynamics (MD) approach. The folding dynamics of heteropolymer chain is studied by MD until the equilibrium globular state, corresponding to free energy minimum, was reached, as in [28]. Equilibrium polymer models are successfully used in polymer modeling of biostructures [31, 35, 36]. Numerical integration of the MD equations is carried out in the OPENMM environment [37, 38].

To determine the coefficients *U*_*attr0ij*_, we developed an iterative algorithm consisting of the following steps: 1. The initial coefficients are equal for all pairs of elements. 2. The ensemble of 1000 conformations is modeled. The initial state of the chain is a self-avoiding polymer coil, the ensemble of final states is determined by folding the chain into a compact condensed state depending on the potentials of volume interactions. 3. A contact map is calculated, where each element (*i, j*) is the frequency of contacts between the subunits *i* and *j*, the calculated contact map is compared with the experimental one, obtained by Hi-C. 4. If the Pearson correlation coefficient between the calculated and experimental maps exceeds 0.97, the iterations are stopped. Otherwise, for each pair (*i, j*), the coefficient *U*_*attr0ij*_ increases, if the estimated frequency of contacts for this pair is lower than the experimental one, and decreases, if higher. 5. Return to step 2. To construct experimental contact maps for chromosomes 12 and 17 in embryonic stem cells hESC and in IMR90 fibroblasts, we use experimental data [9].

The Pearson correlation maps for chromosomes 12 and 17 are obtained similarly to [6]. In short, first, the observed/expected contact matrices are calculated. Then, for each pair (*i, j*), the Pearson correlation coefficients between the *i*-th and *j*-th rows of the “observed/expected contacts” matrix are determined. The only difference from the method [6] is that they calculated the expected contacts throughout the entire genome, and we, within the selected chromosome.

We also use the coarsened model of chromosome 12 with smaller number of monomer types (hereafter referred to as the enlarged model) which is built as follows. First, the potentials *U*_*attr0ij*_ are determined as described above, but for polymer chain of 135 megabase-sized monomers. Second, the monomers are partitioned into several types based on potential similarity by the k-means clustering algorithm. The results of clustering depend on the number of types *n* which was varied from 3 to 9. Here we report the results for *n*=8 for hESC cells and *n*=7 for IMR90 cells which were found to be the minimal numbers allowing to describe the experimental contact maps reasonably well.

## RESULTS

The physical model of a human interphase chromosome reproduces the observed characteristics of Hi-C heatmaps [9] for both pluripotent and differentiated cells with high accuracy (Fig.1 A, B for chromosome 12, Fig.2 A, B for chromosome 17). The maps are normalized to the total number of contacts for all pairs of loci with a genomic distance of more than 100 kbp. Pearson correlation coefficients between calculation and experiment: chromosome 12, hESC cells R = 0.989, IMR90 cells R = 0.992; chromosome 17, hESC R = 0.981, IMR90 R = 0.972. Thus, the model demonstrates the excellent ability of quantitative description of experimental heatmaps for cells in pluripotent and differentiated state.

**Fig. 1.**
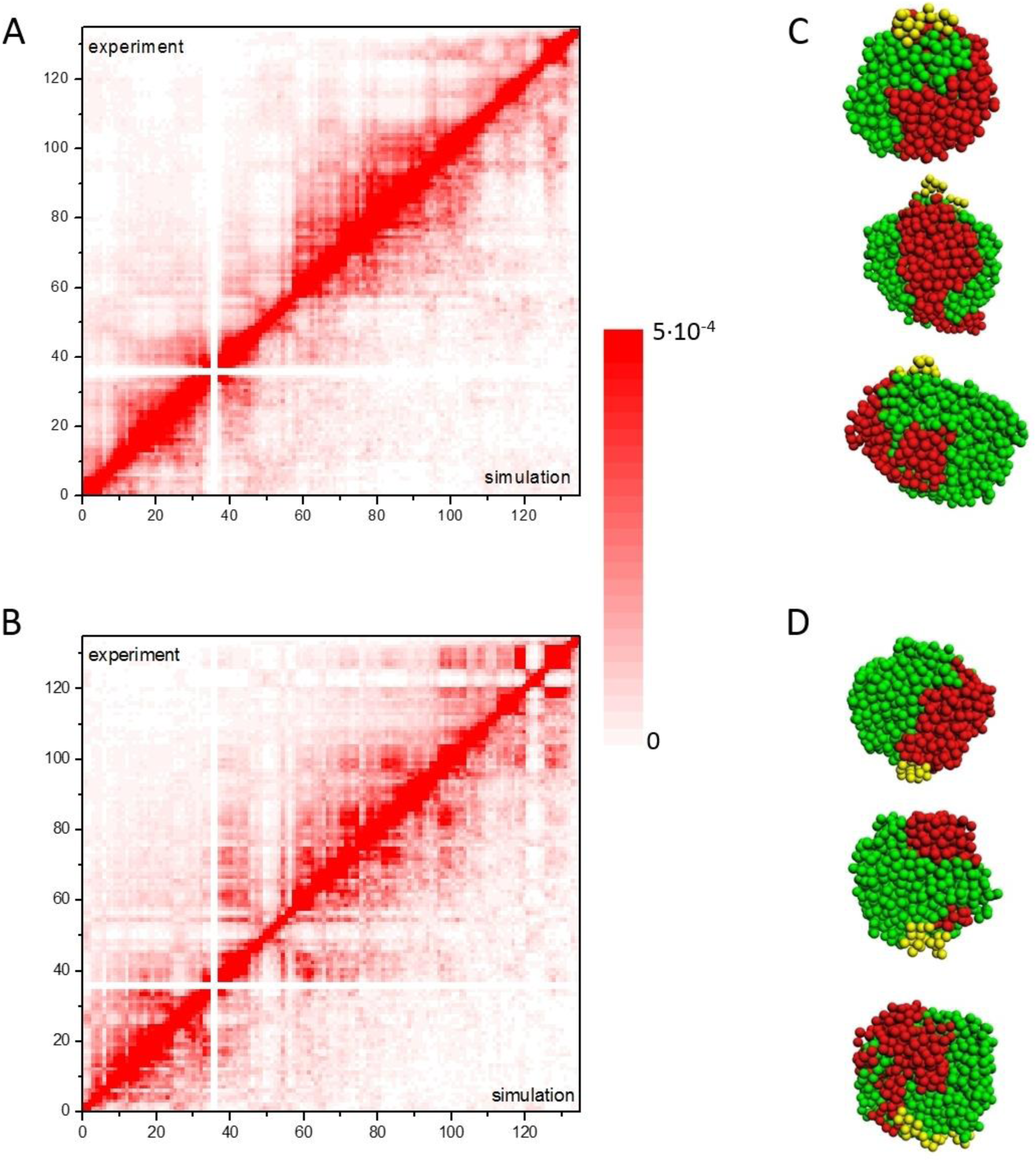
The physical model for chromosome 12 quantitatively describes Hi-C data on human embryonic stem (hESC) and differentiated cells (IMR90). A, C: chromosome 12 in hESC cells. B, D: chromosome 12 in IMR90 cells. A, B: contact maps, predictions by the model for an ensemble of 1000 conformations vs experiment [9]. The genomic distance of the locus from the p-telomere is displayed along the axes. Contacts with participation of the centromeric chromatin (∼35-37 Mbp) are not detected by Hi-C, which corresponds to white crosses. C, D: three conformations from each simulated ensemble. Red subunits, p-arm; green, q-arm; yellow, centromeric region.

**Fig. 2.**
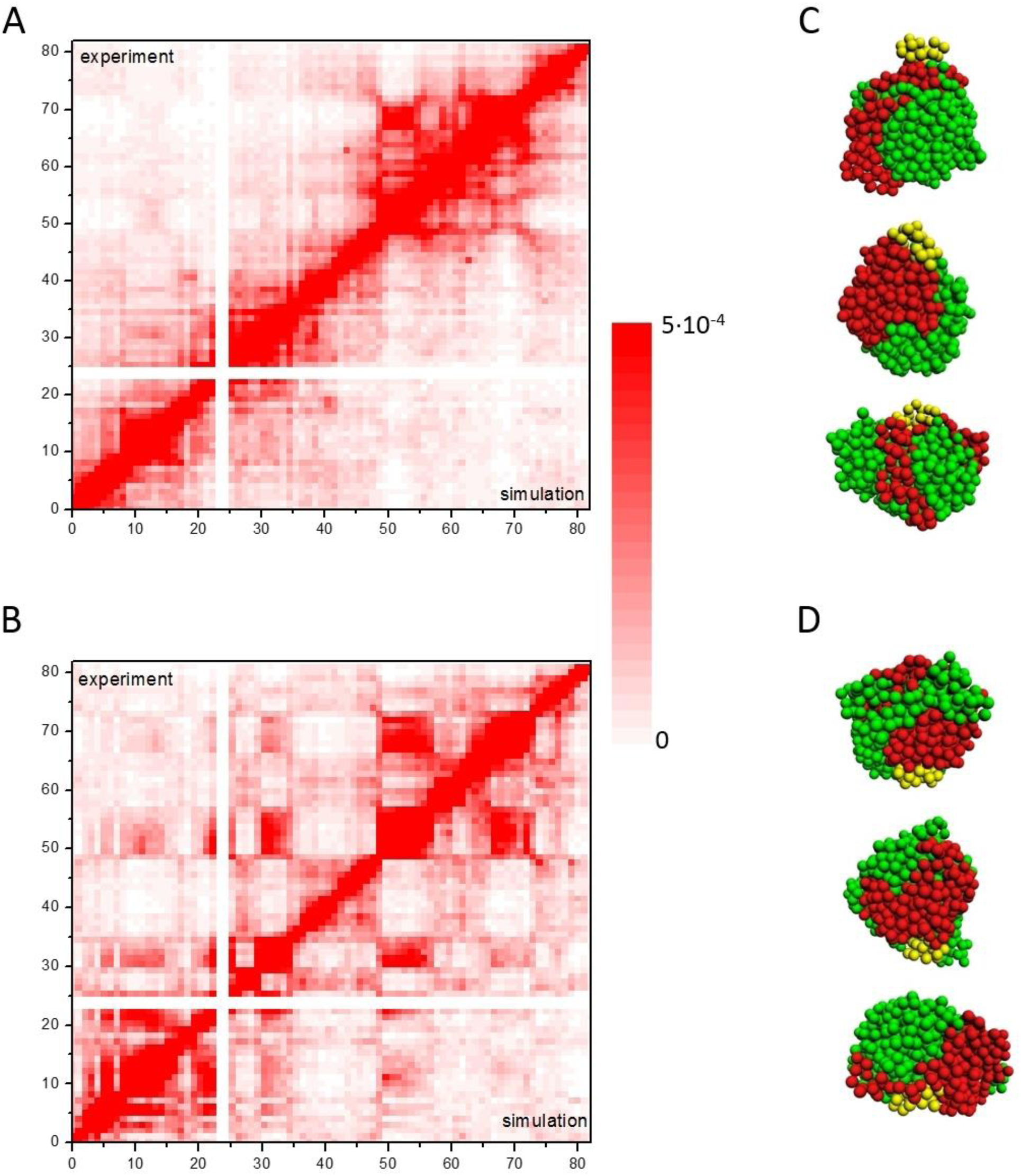
The physical model for chromosome 17 quantitatively describes Hi-C data on human embryonic stem (hESC) and differentiated cells (IMR90). A, C: chromosome 17 in hESC cells. B, D: chromosome 17 in IMR90 cells. A, B: contact maps, predictions by the model for an ensemble of 1000 conformations vs experiment [9]. The genomic distance of the locus from the p-telomere is displayed along the axes. C, D: three conformations from each simulated ensemble. Color indication is the same as in Fig.1 C, D.

The contact maps are calculated for an ensemble of 1000 chromosomal conformations for each cell type. It means that the contact map does not reflect some single conformation. To look at the single-cell distribution of chromosome conformations, three typical conformations from each ensemble are presented in Figs 1 C, D and 2 C, D. Different colors denote chain monomers belonging to different chromosome arms, as well as centromeric regions. The reconstructed chromosome structures for both cell types represent equilibrium heteropolymer globules, as predicted previously [28, 29].

To identify differences in chromosomal contacts specific to the stage of differentiation, we designed two other types of maps. The first type is difference map, where for each pair of loci (*i, j*) the absolute difference in the frequency of contacts is shown between two types of cells for chromosome 12, Fig. 3 A and for chromosome 17, Fig. 3 B. The second type is relative difference map, where for each pair of loci (*i, j*) the difference of contact frequency is divided by the frequency in stem state, hESC cells (chromosome 12, Fig. 3 C; chromosome 17, Fig. 3 D). The brighter the point, the greater the difference, either absolute or relative, between embryonic stem and differentiated cells.

**Fig. 3.**
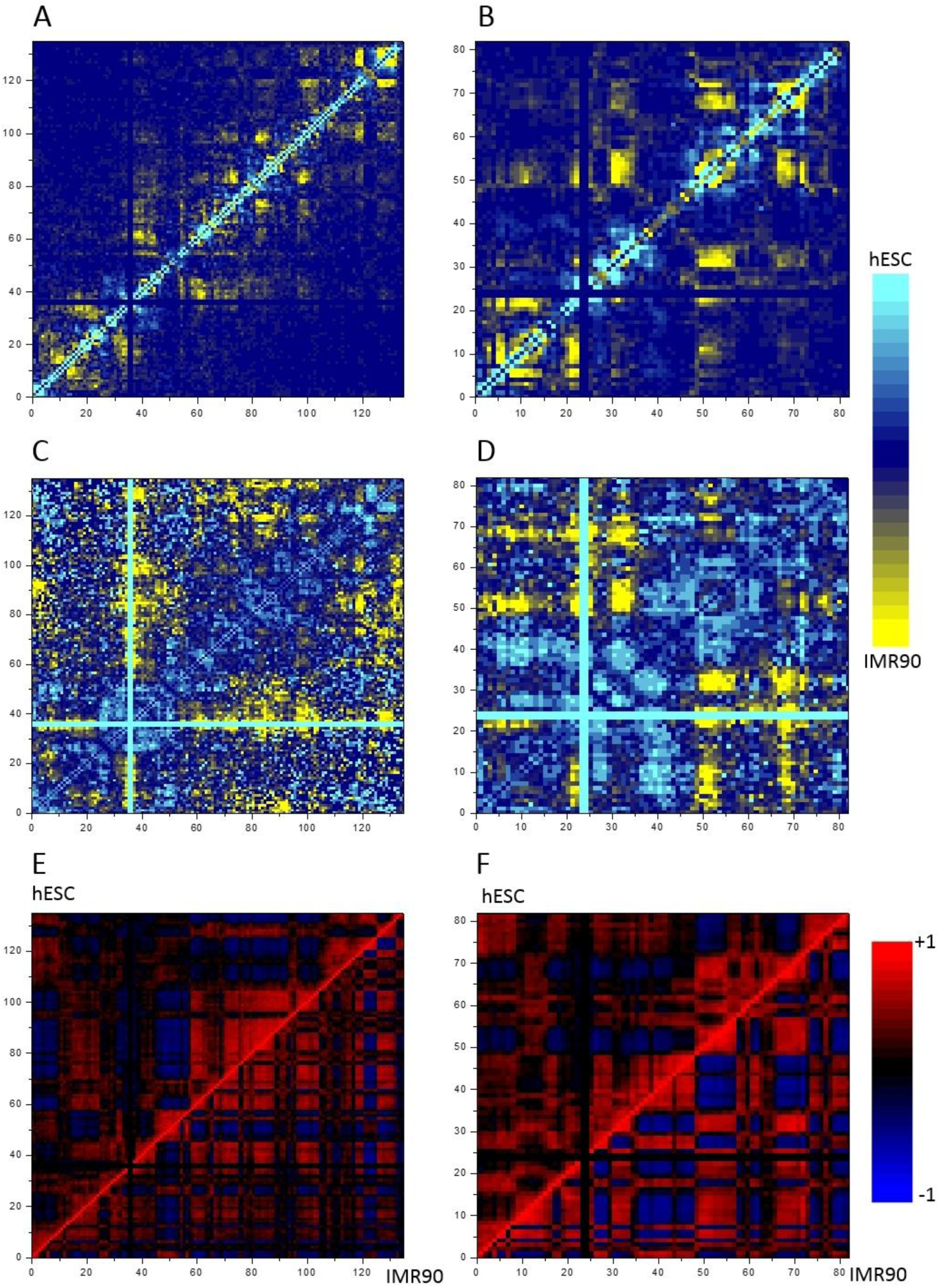
Replacement of compartmentalization of chromosomes 12 and 17 during stem cell differentiation. A, B: the absolute difference maps for contact frequencies in hESC and IMR90 cells. A: chromosome 12, B: chromosome 17. The yellow areas indicate pairs of loci, for which there are more contacts in IMR90, cyan areas indicate pairs of loci, for which there are more contacts in hESC, dark blue areas indicate close numbers of contacts in the two cell types. C, D: the relative difference maps for contact frequencies in hESC and IMR90 cells. C: chromosome 12, D: chromosome 17. The color indication is the same as in panels A, B. E, F: contact correlation maps, comparison of hESC vs IMR90. E: chromosome 12, F; chromosome 17. Colors indicate Pearson correlation coefficients. For detailed explanation see text.

Both map types indicate marked differences between chromosomes/chromosomal contacts in stem and differentiated cells. Map regions with similar differences tend to group together, forming big, about several Mbp, same-color areas on the maps. In certain regions there is abundance of contacts in hESC, in others depletion in hESC and abundance for IMR90. The relative difference maps indicate changes in contact patterns, especially in low-frequency range between elements with large genomic separation (Fig. 3 A-D). Both types of maps clearly demonstrate chromosome-wide replacement of compartmentalization in the process of stem cell differentiation.

Certain chromosome regions demonstrate contact shutdown in the course of differentiation, where dark areas (frequent contacts) on the heatmap are turned into light areas (depleted contacts), meaning drastic decrease of contacts. For example, this can be seen in chromosome 12 for contacts between regions 22-35 Mbp and 37-46 Mbp (Fig.1 A, B, Fig.3 C), and in chromosome 17 for contacts between regions 48-57 Mbp and 58-65 Mbp (Fig.2 A, B, Fig.3 D). Other chromosomal regions demonstrate contacts switching, where redistribution takes place with contacts disappearance in some parts and appearance in other parts of the map. Contact appearance in differentiated cells compared to pluripotent ones is manifested as replacement of the light areas on the contact map with the dark areas. These changes can be seen in chromosome 12 e.g. for contacts between regions 37-47 Mbp and 70-102 Mbp (Fig.1 A, B, Fig.3 C), and in chromosome 17 for contacts between regions 30-35 Mbp and 49-57 Mbp (Fig.2 A, B, Fig.3 D). Thus, structure of chromosomes 12 and 17 demonstrate both shutdown and switching type of compartmentalization replacement in the course of differentiation.

Next, we compare the Pearson correlation maps on the basis of observed-to-expected contact matrix calculations (see Methods) for different cell types, Fig.3 C, D. These maps demonstrate that differentiation is accompanied by increased compartmentalization, which is expressed in more distinct pattern of the grid and in the partitioning of large boxes into several smaller ones. The latter is clearly seen, for example, in the regions 70-95 Mbp and 108-125 Mbp of chromosome 12, as well as in the 0-8 Mbp region of the chromosome 17.

The distribution of the squared gyration radius of both arms of chromosome 12 shows that they are only slightly more condensed in IMR90 cells than in hESC cells (Fig.4 A). For p-arm of chromosome 12, the mean squared gyration radius is 0.931±0.005 μm^2^ in hESC cells and 0.886±0.004 μm^2^ in IMR90 cells, for q-arm 1.296±0.002 and 1.270±0.002 μm^2^, respectively (standard error values are given, for both p- and q-arm the means are significantly different, the Student’s test, p <0.01). We conclude that the chromosome 12 undergoes weak condensation during differentiation. Chromosome 17, on the contrary, has the same level of condensation in both types of cells: for the p-arm, the mean squared gyration radius is 0.682±0.005 μm^2^ in hESC cells and 0.687±0.004 μm^2^ in IMR90 cells, for the q-arm, 0.950±0.003 and 0.957±0.003 μm^2^, respectively (differences in the means for both p- and q-arms are statistically insignificant by the Student’s test, p = 0.4 for the p-arm and 0.055 for the q-arm). The distributions of the squared gyration radius are shown in Fig.4 B.

**Fig. 4.**
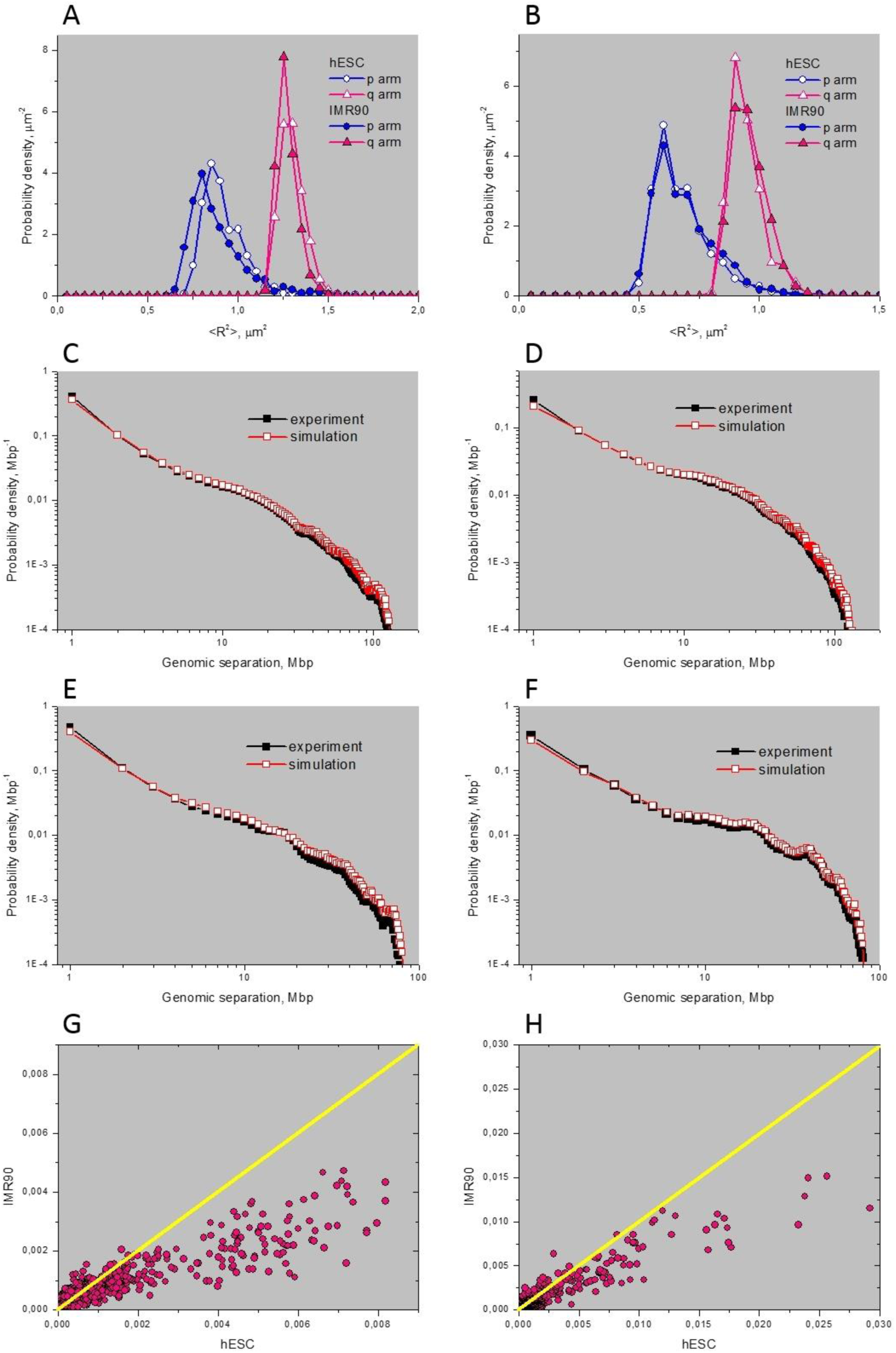
Structural features of chromosomes 12 and 17 during stem cell differentiation. A, B: distribution of the squared gyration radius of the p- and q-arms of human chromosomes for hESC and IMR90 cells. A: chromosome 12, B: chromosome 17. C-F: Distributions of the genomic distance between the contacting loci, simulation *vs* experiment [9]. C, D: chromosome 12; E, F: chromosome 17. C, E: hESC cells; D, F: IMR90 cells. G, H: correlation graphs of the number of contacts in hESC *vs* IMR90 cells. Each point corresponds to one pair of 1 Mbp loci. G: chromosome 12; H: chromosome 17.

To further elucidate the structural differences between the chromosomes in the differentiated and embryonic stem cells, the calculated and experimental distributions of genomic distance between the contacting subunits are compared. For both chromosomes and both cell types the good agreement between simulated and experimental distributions is obtained (Fig.4 C-F). The distributions of genomic distance between the contacting subunits cannot be satisfactorily described by the power law with a single slope. The distributions for both chromosomes and both cell types can be conventionally divided into 4-6 regions characterized by different slopes, Fig.4 C-F. These slopes vary from -0.38 to -3.41. The correlation plot for contact frequencies for different cell types (Fig.4 G, H) further quantifies the differences for both chromosomes. They can reach hundreds percent for some locus pairs.

A distinctive feature of our model is that it does not suggest that the genome/chromosome is partitioned into two [6] or six [12] types of compartments, equivalent to two [39] or six types of polymer blocks of monomers [31]. In the present model, the number of monomer/chromosomal subunits types, equals the total number of monomers divided by the experimental map resolution used (1 Mbp).

The general model without coarsening uses the wide spectrum of chromosomal subunits/chromatin types and, at the same time, is consistent with chromosome compartmentalization seen in the heatmap. To explore conditions when one can lower number of monomer types we simplify the picture according to the following rule. We group different types of chromosomal subunits or chain monomers with resembling volumetric potentials into the same type. For illustration and without loss of accuracy we quantify this enlarged model with 1 Mbp subunits. The number of compartments *n* is varied from 3 to 9. We aim for the “minimal” fit, i.e. minimal number *n* giving the reasonable description of the contact maps for chromosome 12. The minimal fit is obtained for 8 (in hESC) and 7 (in IMR90) compartments.

The coefficients at attractive potentials *U*_*attr0ij*_ (see Methods) depend on the type of interacting elements (*i, j*). The values of these coefficients are given in Tables 1 and 2. Our enlarged model of a heteropolymer chromosome describes the experimental maps with a reasonable, though lower compared to completely heterogeneous non-enlarged model, accuracy. The visual agreement is good, with some exceptions (see Fig.5 A, B). The lower correlation coefficient between experimental and simulated contact maps for the enlarged model is due to the impact of enlargement on the long-range contacts.

**Fig. 5.**
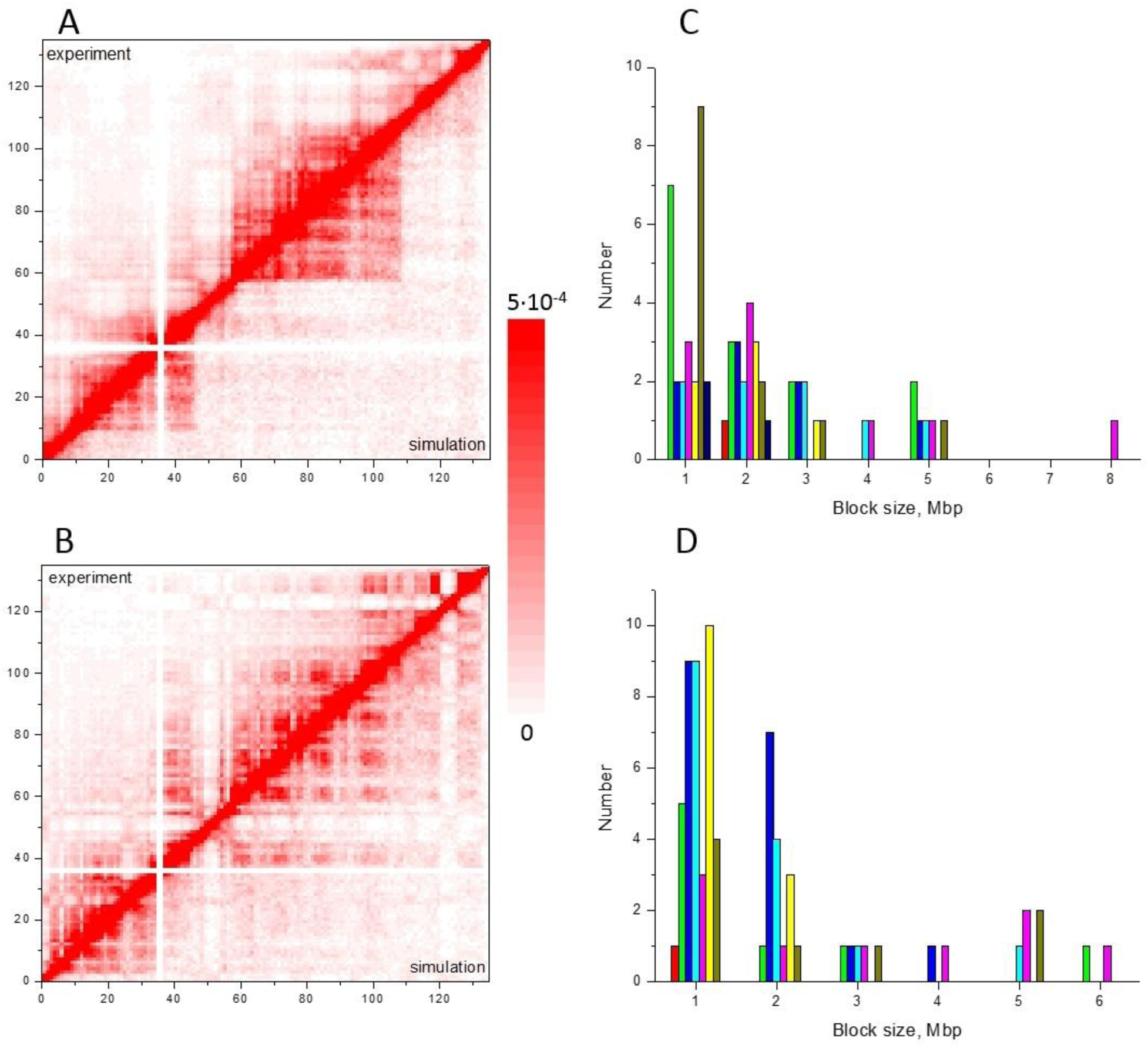
Predictions of enlarged model of chromosome 12 in hESC and IMR90 cells. Calculations were performed for simplification of 1 Mbp monomer size. A, B: contact maps, simulation vs experiment [9]. A: hESC cells. Minimal fit for 8 compartments. Pearson correlation coefficient R=0.929. B: IMR90 cells. Minimal fit for 7 compartments. Pearson correlation coefficient R=0.918. C, D: block size distribution for monomers belonging to different compartment types. C: hESC cells. D: IMR90 cells. Different colors denote different types.

**Table 1.** Values of *U*_*attr0ij*_ (in *kT* units) for subunit pairs (*i, j*) belonging to different compartments, for hESC cells

| type | 1 | 2 | 3 | 4 | 5 | 6 | 7 | 8 |
| --- | --- | --- | --- | --- | --- | --- | --- | --- |
| 1 | 2.398 | 0.925 | 0.964 | 0.936 | 1.138 | 1.092 | 1.000 | 1.191 |
| 2 | 0.925 | 2.171 | 2.390 | 2.093 | 2.429 | 2.553 | 1.934 | 1.491 |
| 3 | 0.964 | 2.390 | 3.490 | 3.356 | 2.523 | 3.946 | 2.033 | 1.683 |
| 4 | 0.936 | 2.093 | 3.356 | 3.864 | 2.550 | 4.295 | 1.500 | 1.884 |
| 5 | 1.138 | 2.429 | 2.527 | 2.550 | 3.268 | 2.948 | 1.995 | 2.545 |
| 6 | 1.092 | 2.553 | 3.946 | 4.295 | 2.948 | 4.350 | 2.024 | 2.085 |
| 7 | 1.000 | 1.934 | 2.033 | 1.500 | 1.995 | 2.024 | 1.930 | 1.205 |
| 8 | 1.191 | 1.491 | 1.683 | 1.884 | 2.545 | 2.085 | 1.205 | 2.524 |

**Table 2.**
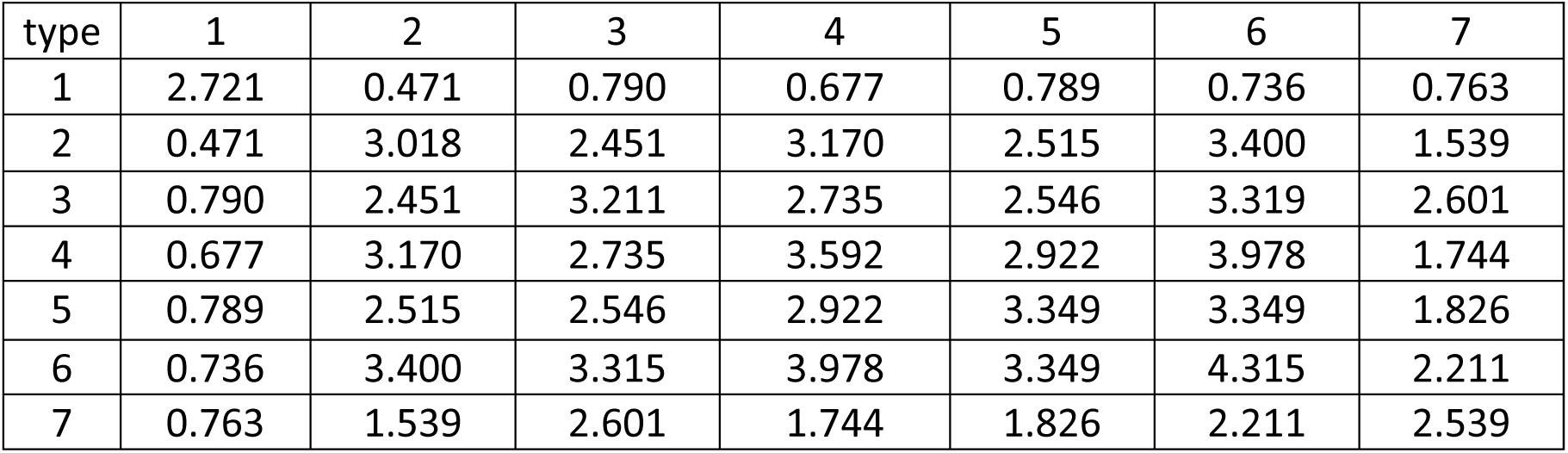
Values of *U*_*attr0ij*_ (in *kT* units) for subunit pairs (*i, j*) belonging to different compartments, for IMR90 cells

| type | 1 | 2 | 3 | 4 | 5 | 6 | 7 |
| --- | --- | --- | --- | --- | --- | --- | --- |
| 1 | 2.721 | 0.471 | 0.790 | 0.677 | 0.789 | 0.736 | 0.763 |
| 2 | 0.471 | 3.018 | 2.451 | 3.170 | 2.515 | 3.400 | 1.539 |
| 3 | 0.790 | 2.451 | 3.211 | 2.735 | 2.546 | 3.319 | 2.601 |
| 4 | 0.677 | 3.170 | 2.735 | 3.592 | 2.922 | 3.978 | 1.744 |
| 5 | 0.789 | 2.515 | 2.546 | 2.922 | 3.349 | 3.349 | 1.826 |
| 6 | 0.736 | 3.400 | 3.315 | 3.978 | 3.349 | 4.315 | 2.211 |
| 7 | 0.763 | 1.539 | 2.601 | 1.744 | 1.826 | 2.211 | 2.539 |

Based on a completely different non-polymeric bioinformatic approach, Rao *et al* found 6 sub-compartments for GM12878 cells [12]. This estimate is surprisingly close to our 7-8 types of compartments deduced from comprehensive polymer chromosome modeling. It reveals that two types of genome compartments, introduced in [6], is not the fundamental characteristics of compartmentalization, but depends on the level of coarsening of the map, sharpness of boundaries, *etc*.

Grouping of monomers by similarity of potentials results in the block-copolymer composition of chromosome chain. The size of monomer blocks varies from 1 to 8 Mbp in hESC cells or from 1 to 6 Mbp in IMR90 cells. Fig.4 C, D shows the distribution of different type monomer blocks according to their size for both cell lines. The distributions are not normalized, along the Y axis the number of blocks of a given size is plotted. The sum of the columns for each compartment is the total number of blocks belonging to given compartment.

## DISCUSSION

The serious problem in the field of physical chromosome modeling is multiple gaps in knowledge of factors responsible for contacts of remote chromosomal elements (see also (29, 38)). By introducing volumetric interactions, the polymer model may capture molecular mechanisms in generalized form, in particular, the concentrations and binding constants of proteins bridging chromatin subunits in proximity. Concentration of bridging proteins may depend on the chromatin type of monomers which is determined by histone modifications, the number of DNAse-I hypersensitive sites, *etc*. Protein-protein and protein-chromatin interactions, which define volumetric interactions, coupled with the principle of minimum free energy, are responsible for establishment of predominant contacts between particular groups of loci on the chromosome responsible for stem cell specification and differentiation.

The areas of heatmap with frequent contacts, manifested as dark rectangles, reflect clusters of chromosome subunits in spatial proximity. We hypothesize that they may serve as units of functional cooperation. The regulatory factors that can reach the given spatial cluster of chromosomal subunits may be the same for all subunits in the cluster. This condition opens up the possibility of coordinated regulation of multiple genes within clusters of remote chromosomal subunits, domains or TADs by the same regulatory factors. This chromosome-wide cooperation may involve chromatin areas of different kinds, not only active-inactive chromatin and genes, but also loci from non-coding chromosome regions.

The model explains the difference in the compartmentalization of pluripotent and terminally differentiated cells by the change in the system of volumetric interactions and type of chromosome/chromatin subunits. How this happens remains unknown, however, one can propose the physical scenario of change of 3D chromosome architecture and the replacement of compartmentalization. Transcription complexes, chromatin-modifying factors, products of gene expression, non-coding RNA and others can be involved as molecular carriers of volumetric interactions, which, together with chromatin type in chromosomal subunits, are changed during the maturation of stem cells. In particular, the products of expression of pluripotent genes and their target genes can modify the higher-order chromatin structure by the change of the types of subunits and cross-linking complexes. These pathways affect the volumetric interactions and can result in compartmentalization alterations. We cannot exclude that developmental genes could be functionally organized according to the compartmentalization rules. Then switching 3D-structures and replacement of compartmentalization would result in changing of functional domains and, consequently, in ordered switching of active/silent genes during differentiation or genome reprogramming and dedifferentiation. In this scenario, the genome structure-function relationships during differentiation are well described by transitions between states, in which the topology of chromosomes (as well as compartmentalization and gene expression profile) on the given stage of differentiation is the consequence of 3D-state on the previous stage. Developmental genes play the role in this process as the switchers which increase the probability of transition from previous to given 3D-structural state. This nonlinear effect is possible owing to the feedback loop between gene products and 3D chromosome structure [41].

In conclusion, the presented polymer model of human interphase chromosomes reproduces the pattern of long-range contacts and compartmentalization in pluripotent and differentiated cells. The model presumes a mechanism that controls the chromosome architecture as a whole and regulates the positions and contacts of remote chromosome loci. Restructuring of chromosomes topology and compartmentalization replacement adjusts the mutual positioning of chromosomal domains and coordinates the expression of large groups of genes during differentiation or genome reprogramming and dedifferentiation.

## ACKNOWLEDGEMENTS

The present work was supported by Russian Foundation for Basic Research grant 14-01-00825 to S.A.

S.A. acknowledges support from the MEPhI Academic Excellence Project (Contract No. 02.a03.21.0005).

